# Age- and sex-related variation in TREC and KREC and the association of rs2204985 with TREC in adults

**DOI:** 10.64898/2026.11.21.768288

**Authors:** Daria V. Shamovskaya, Andrey A. Stepanov, Evgeny A. Khrapov, Svetlana V. Apalko, Ulyana A. Boyarskikh, Maria A. Gordukova, Dmitry A. Kudlay, Maxim L. Filipenko

## Abstract

**Background:** T-cell receptor excision circles (TREC) and kappa-deleting recombination excision circles (KREC) are indirect markers of new T- and B-cell production. Recorded zero values complicate the analysis of age-related and genetic associations in adult cohorts.

**Methods:** We analyzed 574 adult outpatients aged 39–95 years, including 250 men and 324 women. TREC and KREC were quantified by multiplex real-time PCR and normalized per 100,000 cells. The rs2204985 variant was evaluated using age- and sex-adjusted genetic models. A two-part model separated TREC positivity from quantitative TREC levels among participants with positive values. Alternative regressions and receiver operating characteristic analyses were used for sensitivity and age-classification analyses.

**Results:** Age was more strongly associated with log_10_(TREC + 0.1) than with log_10_(KREC + 0.1) (β = −0.0365 and −0.00728 per year, respectively). Women had higher levels in the main-effects models, although the KREC age association differed by sex. There was no clear evidence of an additive rs2204985 association with TREC positivity (odds ratio = 0.836, p = 0.290). Among the 481 TREC-positive participants, the G allele was associated with higher log_10_(TREC) (β = +0.0688 per allele, 95% CI +0.0177 to +0.1199, p = 0.0084; q = 0.0126 across three conditional genetic-model tests). Smearing-corrected marginal predictions for a 58-year-old woman were 40.4 copies/100,000 cells for AA and 53.5 for GG. In the common-positive subset (n = 465), TREC showed higher age-classification AUCs than KREC or their ratio.

**Conclusions:** The rs2204985 association was apparent in conditional TREC levels but not in recorded positivity or full-cohort regression. Analyzing these outcomes separately makes the association easier to interpret, without establishing that zero TREC denotes absent thymopoiesis or that the variant directly alters thymic function.

## 1 Introduction

Immune aging involves several changes involving thymic involution, contraction of the naive T-cell pool, accumulation of terminally differentiated effector and memory cells, and progressive narrowing of lymphocyte receptor repertoires. Thymic output can be assessed indirectly using T-cell receptor excision circles (TREC), circular episomal DNA products generated during TCRA rearrangement in developing alpha-β T cells. Because TREC molecules do not replicate during cell division, their abundance in peripheral blood is enriched in recent thymic emigrants and decreases with both reduced thymic production and peripheral proliferation. The use and interpretation of TREC measurements for TREC as a marker of thymic output has been developed across studies of human aging, adult thymic function, and naive T-cell biology (1-6). Age-related changes in lymphocyte distributions throughout the human body also help explain the value of using TREC as a molecular readout of immune aging (7).

Kappa-deleting recombination excision circles (KREC) serve a similar purpose for the B-cell lineage. KREC are generated during immunoglobulin kappa locus rearrangement and reflect the output of newly formed B cells from bone marrow. Simultaneous TREC/KREC quantification can therefore be used to compare the relative dynamics of T- and B-cell neogenesis, particularly in settings of primary immunodeficiency, immune reconstitution, and population screening (8-10). TREC-based newborn screening for severe combined immunodeficiency (SCID) has become a key public health application of this biology; subsequent triplex approaches incorporating KREC extended the concept to early B-cell maturation defects and broader severe primary immunodeficiencies (11-22).

In adults, TREC and KREC measurements can also help characterize age-related changes in immunity. Thymic epithelial involution begins early in life and continues throughout adulthood, whereas bone-marrow lymphopoiesis is maintained for longer, although it is also reshaped by age, inflammation, and hematopoietic niche remodeling (23-35). Cytomegalovirus exposure is one of the strongest non-genetic modifiers of the peripheral T-cell compartment in older individuals, driving late-differentiated and clonal CD8+ T-cell expansions (36-39). B-cell development and humoral immune responses also change with age, but the pace and mechanisms of age-related decline are not identical to those of T-cell thymopoiesis (40-42). Sex hormones also affect these measurements: gonadal steroids modulate thymic epithelial biology, thymic regeneration, immune reconstitution, and humoral immunity, providing a mechanistic background for sex dimorphism in TREC and KREC phenotypes (43-47).

Genetic differences may also influence TREC levels. The TCRA-TCRD locus is central to V(D)J recombination in thymocytes, and chromatin conformation, enhancer activity, and ordered recombination programs are required for the generation of a diverse T-cell receptor repertoire (48-54). Large-scale studies of immune-cell phenotypes have shown that heritable components contribute to inter-individual variation in immune traits, despite the substantial influence of non-heritable exposures (55-57). A genome-wide association study identified rs2204985 in the TCRA-TCRD region as a reproducible modifier of human thymopoiesis, with the G allele associated with higher TREC levels (58). The same variant has also been investigated in the context of hematopoietic stem cell transplantation (HSCT), where thymic regeneration and donor-related genetic determinants can influence immune recovery and clinical outcomes (59-61).

Analysis becomes difficult when rs2204985 or other genetic modifiers are tested in adult cohorts with wide age ranges. Older individuals frequently have recorded zero TREC values. Such values may reflect low abundance and assay non-detection, but do not establish the absence of thymopoiesis. A two-part approach separates the probability of a recorded positive TREC value from its conditional quantitative level among individuals with TREC > 0. The two analyses describe different features of the data without requiring a biological on/off interpretation of the recorded zeros.

The present study had three aims. First, we described cross-sectional age and sex associations with TREC, KREC, and the TREC/KREC ratio in an adult outpatient cohort aged 39-95 years. Second, we tested whether rs2204985 was associated with TREC using a two-part model distinguishing recorded positivity from quantitative levels among participants with TREC > 0. Third, we compared the discriminatory performance of TREC, KREC, and the TREC/KREC ratio for age-category classification.

## 2 Materials and methods

### 2.1 Study population

The analysis included 574 adults (250 men and 324 women) aged 39–95 years who attended an outpatient clinic for mild acute conditions, such as minor trauma or for preventive examination. The cohort was used as an outpatient approximation of the adult population distribution. Exclusion criteria were active malignancy, immunosuppressive therapy, active immunodeficiency, and the acute phase of infectious disease. All participants provided written informed consent for research use of their biological material. The study was approved by the Ethics Committee of the Institute of Chemical Biology and Fundamental Medicine SB RAS (Protocol No. 1-04 dated 23 April 2025).

### 2.2 Laboratory measurements

Whole blood was collected into EDTA tubes. Genomic DNA was isolated using a standard procedure. TREC and KREC were quantified by multiplex quantitative PCR with simultaneous amplification of a genome-equivalent control and normalization per 100,000 cells, as described previously (22).

The rs2204985 variant in the TCRA-TCRD region was genotyped by real-time PCR using allele-specific fluorescent hybridization probes. The following oligonucleotides were used: 2204985-U, 5′-CTGGCCTGAATTTAGCAACTGG-3′; 2204985-R, 5′-CAGTTTACAAAGGGTTTTCATGTAA-3′; 2204985-PA, HEX-TGTCCTCTCCATTGACCTCC-BHQ2; and 2204985-PG, ROX-TGTCCCCTCCATTGACCTCC-BHQ2. PCR was performed in a final volume of 25 μL containing 65 mM Tris-HCl (pH 8.9), 16 mM ammonium sulfate, 3.0 mM MgCl_2_, 0.05% Tween 20, 0.2 mM dNTPs, 0.45 μM each primer, 0.15 μM each probe, 20-100 ng genomic DNA, and 2 U Taq DNA polymerase. Cycling was performed on a CFX96 instrument (Bio-Rad, USA) with initial denaturation at 96°C for 2 min, followed by 42 cycles of 96°C for 10 s and 60°C for 30 s. Genotypes were called using the instrument software.

### 2.3 Statistical analysis

TREC and KREC distributions were right-skewed and included recorded zero values. Baseline age and sex associations were estimated by ordinary least squares (OLS) using log_10_(TREC + 0.1) and log_10_(KREC + 0.1) in all 574 participants. The log_10_(TREC/KREC) analysis was restricted to the 465 participants with both markers above zero. These main-effects models included age in years and sex (0, men; 1, women), without genotype. Age-by-sex interactions were examined separately. The two-part TREC model comprised maximum-likelihood logistic regression of P(TREC > 0) in the full cohort and OLS regression of log_10_(TREC) among the 481 TREC-positive participants. Both components included age, sex, and the genetic predictor under examination. Here, detectable or positive TREC refers to a recorded value above zero; this definition does not establish an analytical limit of detection or distinguish technical non-detection from biological causes of low TREC. An age-only logistic model was also fitted to describe the association between age and recorded TREC positivity.

The rs2204985 variant was assessed under additive (G-allele dosage 0, 1, or 2), dominant (AG/GG vs. AA), and recessive (GG vs. AA/AG) models. OLS inference used conventional residual-variance standard errors and Student t tests; HC3 heteroscedasticity-robust standard errors were examined as a sensitivity analysis. Logistic regression used inverse-observed-information standard errors and Wald z tests. The three conditional genetic-model tests formed one exploratory family for Benjamini–Hochberg adjustment; q values refer only to that family, and other p values are nominal. Fisher’s combined probability test summarized the two hurdle-component p values for each genetic coding as an exploratory analysis. Marginal arithmetic-mean TREC predictions were calculated as P(TREC > 0) × 10^η^ × mean(10^e^), where η is the conditional OLS predictor and e denotes residuals from that conditional model. The common residual smearing factor corrects retransformation from the log_10_ scale. Predictions were evaluated for a 58-year-old woman; confidence intervals for these marginal predictions were not calculated. Figure 2A uses the conditional model with genotype fixed at AG. Figure 2B compares full-cohort main-effects predictions with genotype fixed at AG and sex standardized to the cohort proportion of women (324/574). The model assumes the same age slope for men and women, rather than estimating a separate slope for each sex.

Sensitivity analyses used full-cohort OLS, Gaussian Tobit regression, and median quantile regression of log_10_(TREC + 0.1), each adjusted for age, sex, and G-allele dosage. Tobit censoring at −1 was an assumption for recorded zeros, not a measured assay detection limit; inference used the observed-information Hessian. Median regression was fitted by linear programming, with standard errors from 2,000 nonparametric pairs-bootstrap resamples (seed 20261121), percentile confidence intervals, and approximate normal Wald p values.

For age-category classification (≥65 and ≥75 years), ROC analyses used the same 465 common-positive participants and scores of −log_10_(TREC), −log_10_(KREC), and −log_10_(TREC/KREC), so larger scores predicted older age. AUC and paired-difference confidence intervals were estimated using 10,000 paired bootstrap resamples stratified by age category (seed 20261121); paired comparisons used two-sided DeLong tests. Detection proportions were summarized in non-overlapping age intervals [39,50), [50,60), [60,70), [70,80), and [80,96), with Wilson 95% confidence intervals.

Hardy–Weinberg equilibrium was assessed by Pearson’s χ^2^ test with one degree of freedom. Tests were two-sided and confidence intervals were 95%. Calculations used Python 3.12.14, NumPy 2.3.5, SciPy 1.17.0, pandas 2.2.3, and scikit-learn 1.8.0. The regression, resampling, ROC, and multiplicity procedures follow established statistical approaches (62–68). Arithmetic-mean retransformation used Duan’s smearing method (69), and binomial proportion intervals used Wilson’s method (70).

## 3 Results

### 3.1 Cohort characteristics

Baseline characteristics are shown in Table 1. The median age was 57.5 years and the mean was 58.4 ± 13.8 years. The non-overlapping age groups contained 200 participants aged 39–49 years (34.8%), 107 aged 50–59 (18.6%), 128 aged 60–69 (22.3%), 96 aged 70–79 (16.7%), and 43 aged 80–95 (7.5%). The male-to-female ratio was 1:1.30. The rs2204985 genotype counts were 149 AA, 271 AG, and 154 GG; allele frequencies were A = 0.4956 and G = 0.5044. The genotype distribution did not depart significantly from Hardy–Weinberg equilibrium (χ^2^ = 1.779, df = 1, p = 0.182).

**Table 1.** Baseline characteristics of the cohort by sex.

| **Parameter** | **Men (n = 250)** | **Women (n = 324)** |
|---|---|---|
| Age, years, mean ± SD (median; range) | 58.9 ± 13.3 (60; 39–88) | 58.0 ± 14.2 (55.5; 39–95) |
| rs2204985 genotype, AA/AG/GG, n (%) | 63 / 121 / 66 (25.2/48.4/26.4) | 86 / 150 / 88 (26.5/46.3/27.2) |
| TREC, copies/10<sup>5</sup> cells, median (Q1–Q3) | 19.1 (5.5–46.5) | 31.4 (10.2–66.5) |
| Recorded zero TREC (TREC = 0), n (%) | 48 (19.2) | 45 (13.9) |
| KREC, copies/10<sup>5</sup> cells, median (Q1–Q3) | 77.3 (34.0–178.2) | 121.6 (60.7–239.1) |
| Recorded zero KREC (KREC = 0), n (%) | 14 (5.6) | 8 (2.5) |
| Both TREC and KREC > 0, n (%) | 194 (77.6) | 271 (83.6) |
| log<sub>10</sub>(TREC/KREC), median (Q1–Q3) | −0.52 (−0.87–−0.22) | −0.54 (−0.89–−0.21) |

The full cohort comprised 574 participants. SD, standard deviation; Q1–Q3, first and third quartiles. Percentages use the sex-specific denominator. TREC and KREC summaries include recorded zeros; log<sub>10</sub>(TREC/KREC) summaries include only participants with both markers > 0 (194 men and 271 women; total n = 465). There were 481 participants with TREC > 0. A recorded zero does not independently establish absence of lymphopoiesis or a validated analytical detection limit. Concentration units follow the assay normalization described in Methods.

### 3.2 Age and sex associations with TREC, KREC, and the TREC/KREC ratio

In the full-cohort age- and sex-adjusted model, log_10_(TREC + 0.1) was negatively associated with age (β = −0.0365 per year, 95% CI −0.0417 to −0.0313, p = 1.86 × 10^−37^). On the shifted marker scale, the fitted association corresponds to approximately 8.1% lower TREC + 0.1 for each additional year of age. Women had higher age-adjusted values (β = +0.2359, p = 0.00145), corresponding to a 1.72-fold difference on that scale. The age-by-sex interaction did not reach the conventional 0.05 threshold (p = 0.0613); estimated slopes were −0.0426 log_10_ units per year in men and −0.0324 in women (Figure 1A). Because participants were measured only once, these estimates do not show how TREC changes within the same person over time.

**Figure 1.**
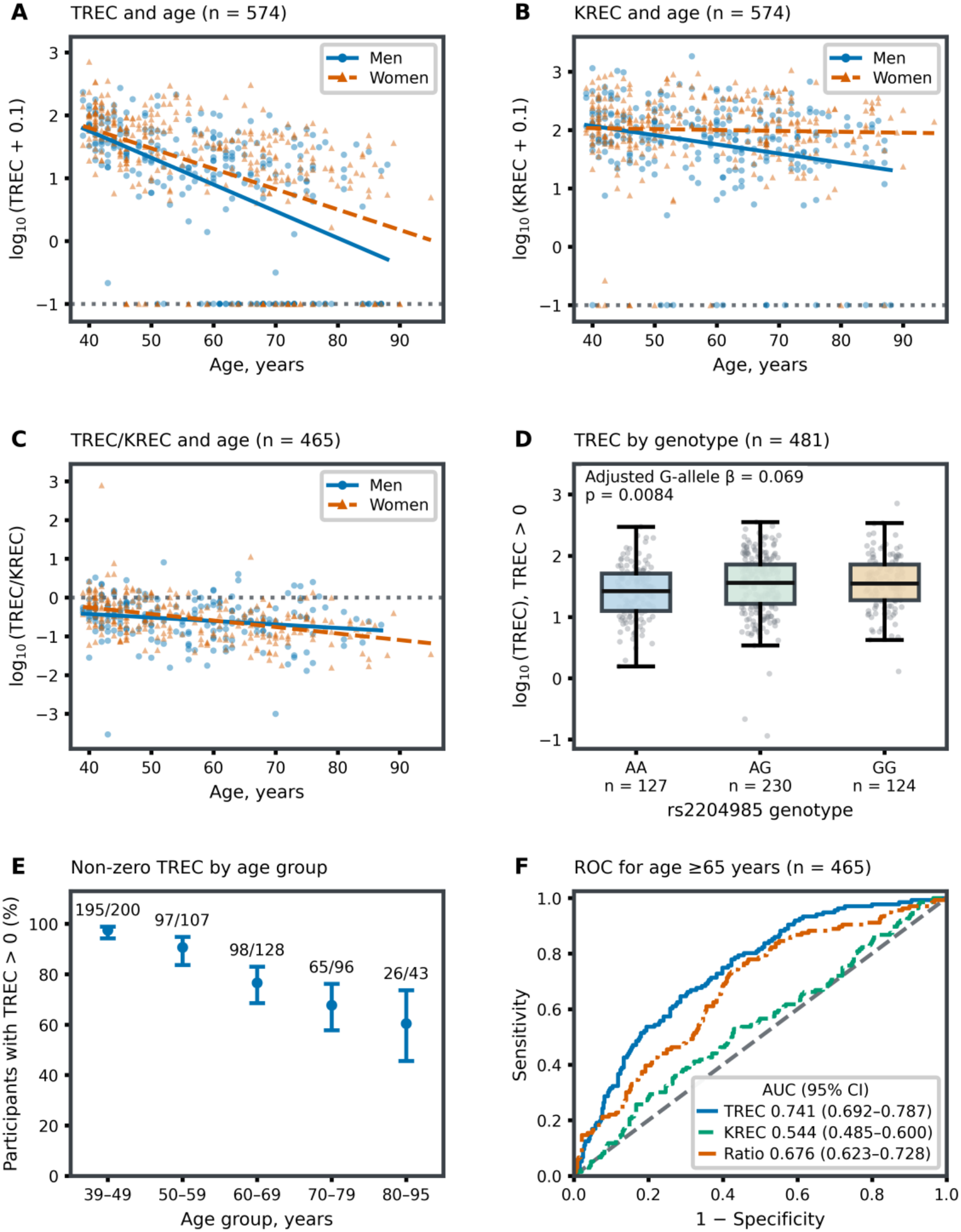
Age, sex, and rs2204985 associations with TREC and KREC. **(A, B)** log_10_(TREC + 0.1) and log_10_(KREC + 0.1) versus age in all 574 participants (250 men, 324 women). Blue circles and solid lines represent men; orange triangles and dashed lines represent women. Lines are separate unadjusted OLS fits by sex and span the observed age range for each sex. Dotted horizontal lines at −1 mark transformed recorded zeros, not a measured assay detection limit. **(C)** log_10_(TREC/KREC) in the 465 participants with both markers > 0 (194 men, 271 women); the dotted line at zero denotes a ratio of 1. **(D)** log_10_(TREC) by genotype among 481 participants with TREC > 0: AA, n = 127; AG, n = 230; GG, n = 124. Boxes show the median and first–third quartiles; whiskers extend to the most extreme observations within 1.5 interquartile ranges. All observations are shown once with horizontal jitter, including points beyond the whiskers. The annotation reports the age- and sex-adjusted additive G-allele coefficient; the boxes themselves are unadjusted. **(E)** Percentage with TREC > 0 in non-overlapping age groups; points show proportions, bars show Wilson 95% confidence intervals, and labels give positive/total counts. **(F)** ROC curves for age ≥65 years in the same 465 common-positive participants (136 older, 329 younger), using −log_10_(marker) scores. TREC is blue/solid, KREC green/dashed, and TREC/KREC orange/dash-dot. The diagonal denotes chance discrimination; legend values are AUCs with 95% percentile confidence intervals from 10,000 paired stratified bootstrap resamples. TREC and KREC values use the normalization described in Methods (copies per 10^5^ cells).

The proportion of recorded zero TREC values increased across age groups, from 5/200 (2.5%) at 39–49 years to 17/43 (39.5%) at 80–95 years. The corresponding TREC-positive proportions were 97.5%, 90.7%, 76.6%, 67.7%, and 60.5%; Wilson 95% confidence intervals are shown in Figure 1E. In the age-only logistic model, the age coefficient was −0.0760 (p = 3.29 × 10^−15^), with an odds ratio for TREC positivity of 0.927 per additional year (95% CI 0.910–0.945). The fitted probability reached 50% at approximately 84.6 years, an estimate specific to this cohort and model.

KREC showed a weaker overall age association in the main-effects model (β = −0.00728 for log_10_(KREC + 0.1), 95% CI −0.01161 to −0.00295, p = 0.00101), corresponding to approximately 1.7% lower KREC + 0.1 per additional year of age. Women had higher values in this model (β = +0.2256, p = 0.000252). However, the age-by-sex interaction was supported (β = +0.0142, p = 0.00152): the estimated age slope was −0.0158 in men (p = 4.39 × 10^−5^) and −0.00156 in women (p = 0.548). The pooled slope therefore does not describe the sexes equally (Figure 1B). Figure 2B provides a standardized comparison of the common slopes from the main-effects TREC and KREC models. Recorded zero KREC values occurred in 22/574 participants (3.8%); their proportion was 14.0% in the 80–95-year group (6/43).

**Figure 2.**
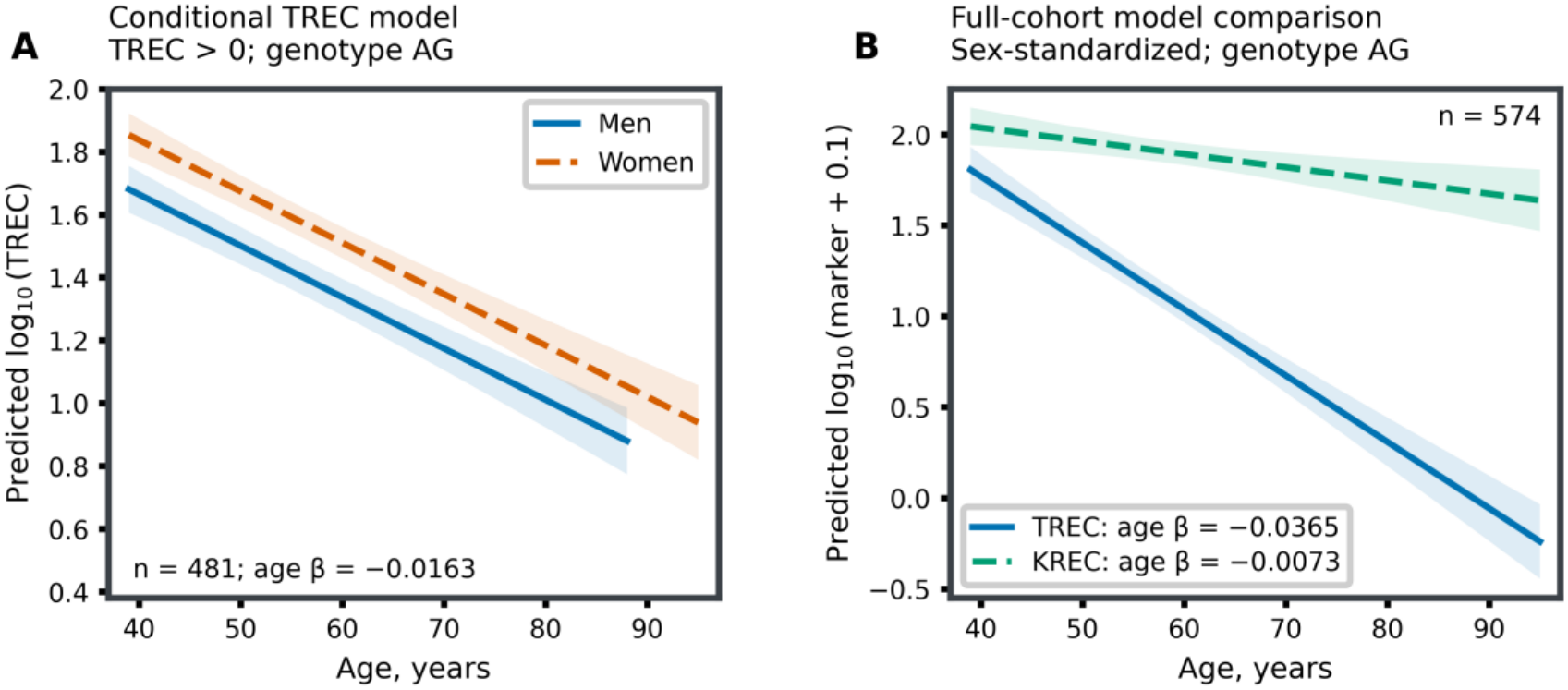
Distinct conditional and full-cohort age models. **(A)** Predictions from log_10_(TREC) ∼ age + sex + G-allele dosage among participants with TREC > 0 (n = 481), with genotype fixed at AG. Men and women are shown separately over their observed age ranges (39–88 and 39–95 years, respectively). The common conditional age coefficient is −0.01635 log_10_ units per year of age. **(B)** Full-cohort models of log_10_(TREC + 0.1) and log_10_(KREC + 0.1) (n = 574), each including age, sex, and G-allele dosage. Genotype is fixed at AG, and predictions are standardized to the cohort sex distribution (324/574 women). The common age coefficients are −0.03652 and −0.00727 log_10_ units for TREC and KREC, respectively. These pooled main-effects predictions do not represent separate sex-specific trajectories; the KREC age-by-sex interaction is examined separately and illustrated in Figure 1B. Shaded bands in both panels show pointwise 95% confidence intervals for the fitted mean on the displayed log scale, not individual prediction intervals. The panels use different populations and transformations. All curves describe cross-sectional age associations, not within-person annual rates. Marker values follow the copies-per-10^5^-cells normalization in Methods.

The TREC/KREC ratio was evaluated in the 465 participants with both markers above zero. In the age- and sex-adjusted model, log_10_(TREC/KREC) decreased with age (β = −0.01395 per year, 95% CI −0.01759 to −0.01031, p = 2.78 × 10^−13^; Figure 1C). The estimated sex coefficient was small and imprecise (β = +0.0440 for women, 95% CI −0.0525 to +0.1404, p = 0.371). The adjusted R^2^ was 0.106 (Table 2). The weak association with sex does not mean that using the ratio eliminates sex-related influences or other confounding.

**Table 2.**
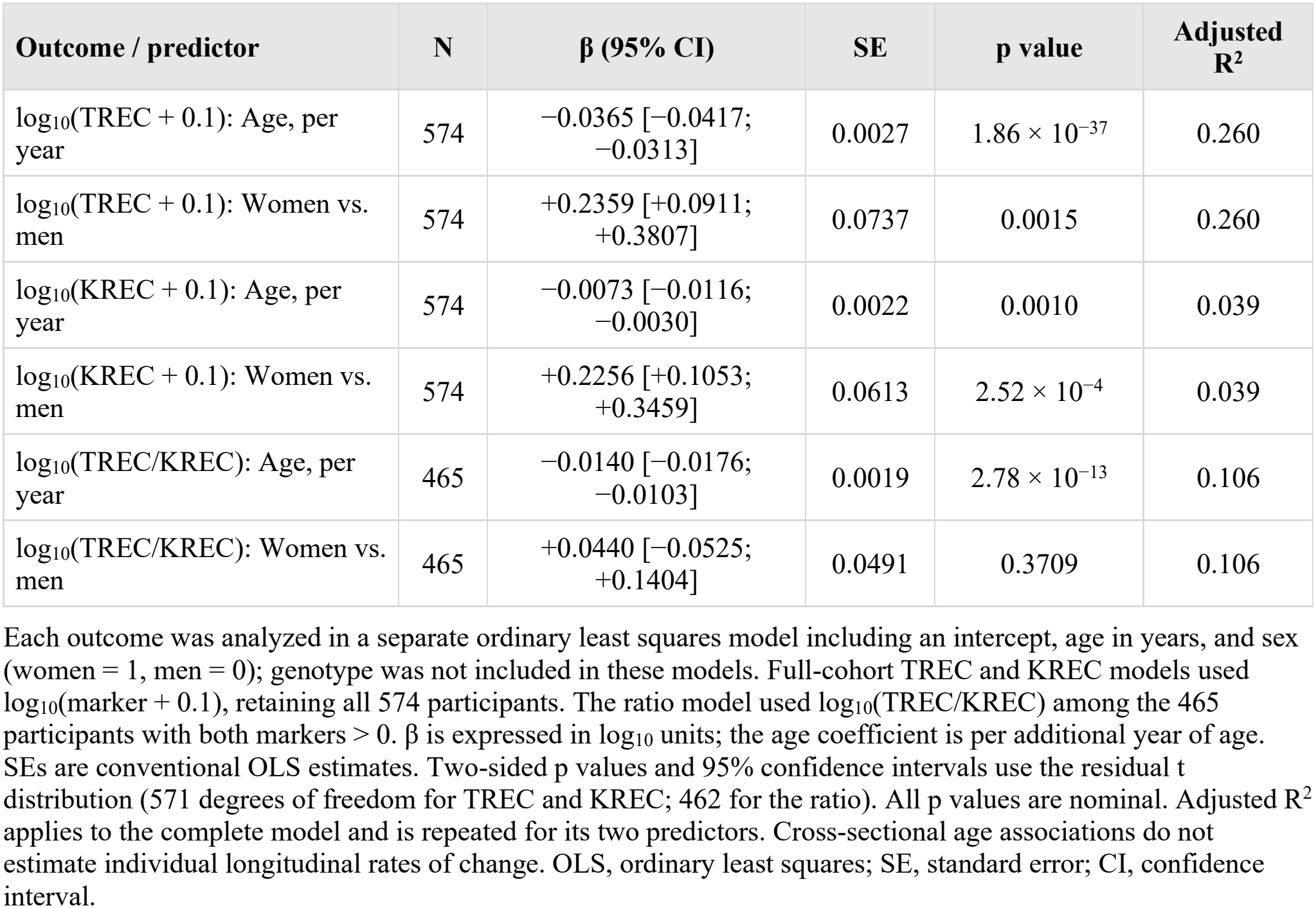
Age- and sex-adjusted associations of TREC, KREC, and the TREC/KREC ratio.

| **Outcome / predictor** | **N** | **β (95% CI)** | **SE** | **p value** | **Adjusted R<sup>2</sup>** |
|---|---|---|---|---|---|
| log<sub>10</sub>(TREC + 0.1): Age, per year | 574 | −0.0365 [−0.0417; −0.0313] | 0.0027 | 1.86 × 10<sup>−37</sup> | 0.260 |
| log<sub>10</sub>(TREC + 0.1): Women vs. men | 574 | +0.2359 [+0.0911; +0.3807] | 0.0737 | 0.0015 | 0.260 |
| log<sub>10</sub>(KREC + 0.1): Age, per year | 574 | −0.0073 [−0.0116; −0.0030] | 0.0022 | 0.0010 | 0.039 |
| log<sub>10</sub>(KREC + 0.1): Women vs. men | 574 | +0.2256 [+0.1053; +0.3459] | 0.0613 | 2.52 × 10<sup>−4</sup> | 0.039 |
| log<sub>10</sub>(TREC/KREC): Age, per year | 465 | −0.0140 [−0.0176; −0.0103] | 0.0019 | 2.78 × 10<sup>−13</sup> | 0.106 |
| log<sub>10</sub>(TREC/KREC): Women vs. men | 465 | +0.0440 [−0.0525; +0.1404] | 0.0491 | 0.3709 | 0.106 |

Each outcome was analyzed in a separate ordinary least squares model including an intercept, age in years, and sex (women = 1, men = 0); genotype was not included in these models. Full-cohort TREC and KREC models used log<sub>10</sub>(marker + 0.1), retaining all 574 participants. The ratio model used log<sub>10</sub>(TREC/KREC) among the 465 participants with both markers > 0. β is expressed in log<sub>10</sub> units; the age coefficient is per additional year of age. SEs are conventional OLS estimates. Two-sided p values and 95% confidence intervals use the residual t distribution (571 degrees of freedom for TREC and KREC; 462 for the ratio). All p values are nominal. Adjusted R<sup>2</sup> applies to the complete model and is repeated for its two predictors. Cross-sectional age associations do not estimate individual longitudinal rates of change. OLS, ordinary least squares; SE, standard error; CI, confidence interval.

### 3.3 rs2204985 associations with recorded TREC positivity and conditional levels

Recorded zero TREC values accounted for 93/574 participants (16.2%), with a larger proportion at older ages. We analyzed the presence of a recorded TREC value separately from its level when TREC was above zero. It did not classify participants as having absent versus active thymopoiesis: the available measurements cannot separate assay-related non-detection, low circulating TREC abundance, and their biological determinants.

There was no clear evidence of an additive rs2204985 association with recorded TREC positivity (β = −0.1794, p = 0.290; OR = 0.836, 95% CI 0.600–1.165). Among TREC-positive participants, the G allele was associated with higher log_10_(TREC) after adjustment for age and sex (β = +0.0688 per allele, 95% CI +0.0177 to +0.1199, p = 0.0084, q = 0.0126), corresponding to a 1.17-fold conditional geometric-mean difference per G allele. The dominant model yielded β = +0.1253 (95% CI +0.0418 to +0.2089, p = 0.0034, q = 0.0101); the recessive estimate was less precise (β = +0.0603, p = 0.163, q = 0.163). The q values apply only to these three conditional genetic tests. HC3 sensitivity p values were 0.0061 for the additive model and 0.0036 for the dominant model. Exploratory Fisher combined p values were 0.0171 and 0.0153 for the additive and dominant models, respectively. In the additive two-part model, smearing-corrected marginal arithmetic-mean predictions for a 58-year-old woman were 40.4, 46.6, and 53.5 copies/100,000 cells for AA, AG, and GG, respectively, a 32.6% higher prediction for GG than AA (Table 3). The genotype distributions and conditional model predictions are shown in Figures 1D and 2A.

**Table 3.** Age- and sex-adjusted rs2204985 associations in the two components of the TREC hurdle model.

| **Genetic model** | **Detection: logit β (95% CI); p** | **Conditional level: β (95% CI); p** | **Combined Fisher p** |
|---|---|---|---|
| Additive (G-allele dosage) | −0.179 [−0.512; +0.153]; p = 0.2897 | +0.069 [+0.018; +0.120]; p = 0.0084 | 0.0171 |
| Dominant (AG/GG vs. AA) | −0.135 [−0.693; +0.423]; p = 0.6350 | +0.125 [+0.042; +0.209]; p = 0.0034 | 0.0153 |
| Recessive (GG vs. AA/AG) | −0.339 [−0.865; +0.186]; p = 0.2054 | +0.060 [−0.025; +0.145]; p = 0.1630 | 0.1472 |

The detection component is logistic regression for P(TREC > 0) in all 574 participants; its β is a log odds coefficient. The conditional component is OLS for log<sub>10</sub>(TREC) in the 481 participants with TREC > 0; its β is in log<sub>10</sub> units. Every model includes age and sex. Genotype coding was AA/AG/GG = 0/1/2 for the additive model, AA = 0 and AG/GG = 1 for the dominant model, and AA/AG = 0 and GG = 1 for the recessive model. Logistic intervals and p values use Wald z inference; conditional OLS uses t inference with 477 degrees of freedom. Fisher's method combines the two nominal component p values using a χ² reference distribution with four degrees of freedom. For a 58-year-old woman, additive-model marginal arithmetic mean TREC values are AA 40.4, AG 46.6, and GG 53.5 copies/10<sup>5</sup> cells (GG vs. AA: +32.6%). These were computed as P(TREC > 0) × 10<sup>η</sup> × 1.47499; η is the fitted conditional log<sub>10</sub>(TREC), and the final multiplier is the Duan residual smearing factor. Multiplying detection probability by the conditional geometric mean alone does not give the marginal arithmetic mean. Non-significance in the detection component does not establish absence of a genotype effect on detection. Benjamini–Hochberg correction was applied only to the three conditional genetic tests (additive, dominant, and recessive): q = 0.0126, 0.0101, 0.1630, respectively. Detection-component, combined Fisher, and other exploratory p values remain nominal.

The age- and sex-adjusted additive genotype estimate differed across the analyzed outcomes (Table 4). Full-cohort OLS of log_10_(TREC + 0.1) gave β = −0.0045 (p = 0.928), and the Tobit sensitivity model gave β = −0.0167 (p = 0.780). Median quantile regression gave β = +0.0268 (bootstrap SE = 0.0370, approximate Wald p = 0.470; percentile 95% CI −0.0380 to +0.1090). The conditional OLS component gave β = +0.0688 (p = 0.0084). The models answer somewhat different questions and treat recorded zeros differently. Their comparison supports reporting detection and conditional-level associations separately; a significant result in one model does not make that model more biologically valid.

**Table 4.** Additive rs2204985 associations across four analytical approaches to TREC.

| **Method, outcome, and population** | **N** | **β (95% CI)** | **SE** | **p value** |
|---|---|---|---|---|
| OLS: log<sub>10</sub>(TREC + 0.1), all participants | 574 | −0.005 [−0.103; +0.094] | 0.050 | 0.9284 |
| Tobit: log<sub>10</sub>(TREC + 0.1), all participants | 574 | −0.017 [−0.134; +0.100] | 0.060 | 0.7797 |
| Median regression: log<sub>10</sub>(TREC + 0.1), all participants | 574 | +0.027 [−0.038; +0.109] | 0.037 | 0.4699 |
| Hurdle detection: logit[P(TREC > 0)], all participants | 574 | −0.179 [−0.512; +0.153] | 0.169 | 0.2897 |
| Hurdle conditional level: log<sub>10</sub>(TREC), TREC > 0 | 481 | +0.069 [+0.018; +0.120] | 0.026 | 0.0084 |

The four approaches are full-cohort OLS, Tobit regression, median quantile regression, and the two-component hurdle model; the hurdle components are shown separately. All models include age, sex, and G-allele dosage (AA/AG/GG = 0/1/2). For OLS, SEs are conventional and two-sided p values and CIs use t inference. Tobit assumes a Gaussian latent outcome with left censoring at log<sub>10</sub>(0.1) = −1; SEs are obtained from the inverse observed-information Hessian, with Wald z p values and CIs. This censoring threshold is a modeling assumption for recorded zeros, not an independently established assay detection limit. Median regression was fitted by linear programming. Its SE is the standard deviation across 2,000 nonparametric pairs-bootstrap replicates (seed 20261121); its CI is the 2.5th–97.5th bootstrap percentile interval, and its p value is an approximate two-sided normal Wald test using β/SE. The logistic hurdle component uses maximum-likelihood SEs and Wald z inference; the conditional component uses OLS t inference. All p values are nominal. Coefficients have different meanings across outcomes and populations; differences in their significance do not by themselves establish superiority of one approach. SE, standard error; CI, confidence interval.

In the age- and sex-adjusted full-cohort KREC model, the additive rs2204985 coefficient was +0.0118 for log_10_(KREC + 0.1) (95% CI −0.0703 to +0.0939, p = 0.778). Absolute lymphocyte counts were not included in the analytical dataset and were not evaluated in this analysis. A non-significant KREC result is not enough to conclude that the variant acts specifically in the T-cell lineage.

### 3.4 Discrimination of age categories by TREC, KREC, and the TREC/KREC ratio

ROC analyses used the same 465 participants with positive TREC and KREC for all three markers. There were 136 participants aged ≥65 years and 50 aged ≥75 years. TREC had an AUC of 0.741 for age ≥65 years (paired stratified bootstrap 95% CI 0.692–0.787) and 0.742 for age ≥75 years (95% CI 0.678–0.804). KREC had weaker discrimination, with AUCs of 0.544 (95% CI 0.485–0.600) and 0.560 (95% CI 0.480–0.638), respectively (Table 5; Figure 1F).

**Table 5.** Discrimination of age categories by TREC, KREC, and the TREC/KREC ratio in a common positive-marker subset.

| **Marker** | **AUC for ≥65 years** | **95% bootstrap CI** | **AUC for ≥75 years** | **95% bootstrap CI** |
|---|---|---|---|---|
| TREC | 0.741 | [0.692; 0.787] | 0.742 | [0.678; 0.804] |
| KREC | 0.544 | [0.485; 0.600] | 0.560 | [0.480; 0.638] |
| TREC/KREC | 0.676 | [0.623; 0.728] | 0.658 | [0.580; 0.733] |

All markers were evaluated in the same 465 participants with TREC > 0 and KREC > 0. The ≥65-year analysis comprised 136 older and 329 younger participants; the ≥75-year analysis comprised 50 older and 415 younger participants. Scores were −log<sub>10</sub>(TREC), −log<sub>10</sub>(KREC), and −log<sub>10</sub>(TREC/KREC), so higher scores predicted older age. Confidence intervals are 2.5th–97.5th percentiles of 10,000 paired bootstrap resamples stratified by age-category outcome (seed 20261121). Paired contrasts use two-sided DeLong tests; all p values below are nominal. ΔAUC is calculated from unrounded estimates as the first marker minus the second. At age ≥65 years, TREC/KREC − TREC: ΔAUC = −0.0643, p = 0.0161; KREC − TREC: ΔAUC = −0.1970, p = 5.10 × 10<sup>−10</sup>; TREC/KREC − KREC: ΔAUC = +0.1326, p = 0.0083. At age ≥75 years, TREC/KREC − TREC: ΔAUC = −0.0842, p = 0.0234; KREC − TREC: ΔAUC = −0.1822, p = 5.91 × 10<sup>−5</sup>; TREC/KREC − KREC: ΔAUC = +0.0979, p = 0.1742. These are within-sample age-classification results and do not establish clinical diagnostic performance.
AUC, area under the receiver operating characteristic curve; CI, confidence interval.

The TREC/KREC ratio had lower AUCs than TREC alone. For age ≥65 years, its AUC was 0.676 (95% CI 0.623–0.728), and the paired difference relative to TREC was −0.0643 (bootstrap 95% CI −0.1174 to −0.0124; paired DeLong p = 0.0161). For age ≥75 years, the ratio AUC was 0.658 (95% CI 0.580–0.733), with a difference of −0.0842 (bootstrap 95% CI −0.1586 to −0.0138; paired DeLong p = 0.0234). In these exploratory analyses, the ratio classified age groups no better than TREC alone in the common-positive subset; they do not establish the relative clinical prognostic value of these markers.

## 4 Discussion

This study analyzed TREC and KREC as molecular readouts related to lymphopoiesis in 574 adult outpatients and evaluated the rs2204985 association with TREC using a two-part model. The model distinguished recorded TREC positivity from quantitative levels among participants with positive values. The main results are discussed below.

First, TREC showed a stronger inverse association with age than KREC in the pooled age- and sex-adjusted analyses, and recorded zero values were more frequent for TREC. This is consistent with the known decline in thymic activity with age, but TREC abundance in blood also reflects peripheral cell division and cell composition (1–6). The slopes describe between-person differences on the log_10_(marker + 0.1) scale; they do not estimate annual within-person loss or directly compare rates of T- and B-cell production.

Second, women had higher age-adjusted TREC values. KREC showed an exploratory age-by-sex interaction (nominal p = 0.0015), with a more negative estimated age slope in men and a near-flat slope in women. The overall KREC estimate therefore masks a difference between men and women. Similar differences between men and women have been reported in earlier studies of lymphopoiesis (43–47) but do not identify a hormonal mechanism. The small, imprecise sex coefficient for TREC/KREC likewise does not establish that the ratio removes shared sex-related influences.

Third, TREC/KREC did not improve age-category discrimination over TREC alone in the 465 participants with both markers positive. The ratio had a lower AUC at the two age cutoffs in this subgroup; it does not show that KREC contributes only noise or exclude other research uses of the ratio. These comparisons, made within a single cohort, cannot demonstrate clinical usefulness or independently validated performance as a marker of thymic aging.

The rs2204985 G allele was associated with higher log_10_(TREC) among the 481 participants with positive TREC values. Neither the recorded-positivity model nor the full-cohort log_10_(TREC + 0.1) model provided clear evidence of a genotype association. The conditional and full-cohort analyses address different outcomes and populations, so their differing results do not establish that the genetic association is biologically restricted to active thymopoiesis. Because rs2204985 lies in the TCRA-TCRD region, its possible functional effects warrant investigation (48–54, 58), but these data do not identify how the variant affects TREC abundance or demonstrate a thymic on/off state.

Although the rs2204985 estimate for KREC did not differ significantly from zero, its confidence interval included effects in both directions. We cannot conclude from this result that the association is specific to T cells; population structure, measurement effects, and residual confounding remain possible explanations. Further studies should test whether the conditional TREC association reflects a locus-specific mechanism.

A two-part model allows recorded positivity and TREC levels above zero to be analyzed separately when zero values are common. Its conditional genotype estimate should be reported alongside the positivity component and relevant full-cohort analyses. OLS, Tobit, and median regression use distinct assumptions and target quantities; their p-values alone do not establish superior power or biological validity of the two-part approach. The association among TREC-positive participants needs to be confirmed in another cohort, with careful attention to assay performance.

### 4.1 Limitations

The study has several limitations. The cohort was outpatient-based and not a random population sample; latent comorbidities and healthcare-seeking behavior may therefore have influenced marker distributions. Individuals younger than 39 years were not included. Data on cytomegalovirus serostatus, body mass index, smoking, C-reactive protein, medication exposures other than exclusionary immunosuppression, and additional inflammatory confounders were unavailable. The cross-sectional design does not permit estimation of individual longitudinal rates of decline. Flow-cytometric data on naive and memory T- and B-cell subsets were unavailable, limiting presented interpretation. Assay detection and quantification limits and the reasons for recorded zeros were not documented in the available data, so zero TREC cannot establish cessation of thymopoiesis. The conditional genotype association applies to the TREC-positive subset and requires replication; an age gradient in recorded positivity does not validate a separate biological on/off process. The exploratory KREC age-by-sex interaction also requires confirmation. Finally, ROC comparisons were restricted to participants with both markers positive and were not independently validated.

### 4.2 Conclusion

In 574 adult outpatients aged 39-95 years, TREC showed a stronger inverse cross-sectional association with age than KREC in pooled analyses. Women had higher age-adjusted TREC, while KREC age patterns differed by sex. TREC/KREC did not improve discrimination of older age over TREC alone in participants with both markers positive. The rs2204985 G allele was associated with higher TREC among participants with positive values, but evidence of an association was not detected for recorded positivity or full-cohort log_10_(TREC + 0.1). The two-part analysis shows why these results should be reported separately. The association observed among TREC-positive participants warrants further study without establishing causality, annual within-person decline, or biological cessation of thymopoiesis in participants with recorded zeros.

## Data availability statement

The datasets presented in this article are not readily available because they contain individual-level clinical and genetic data. Requests to access anonymized data should be directed to the corresponding author, Maxim L. Filipenko, and will be considered in accordance with the informed-consent conditions and applicable institutional requirements.

## Ethics statement

The studies involving human participants were reviewed and approved by the Ethics Committee of the Institute of Chemical Biology and Fundamental Medicine SB RAS (Protocol No. 1-04 dated 23 April 2025). The studies were conducted in accordance with local legislation and institutional requirements. Written informed consent was obtained from all participants for the research use of biological material.

## Author contributions

DVS and EAK performed DNA extraction and molecular measurements. AAS collected clinical material. UAB and MAG performed data analysis. SVA and DAK edited the manuscript. MLF conceived the study, analyzed data, and wrote and edited the manuscript. All authors contributed to the article and approved the submitted version.

## Acknowledgments

The authors acknowledge the participants who provided biological material for this study. During manuscript preparation, OpenAI ChatGPT (GPT-5.5 Thinking; OpenAI, San Francisco, CA, USA) was used for translation into English and editorial language polishing. The authors are responsible for verification of the analyses, interpretation, references, and final submitted version.

## Abbreviations

AUC: area under the receiver operating characteristic curve
CI: confidence interval
FDR: false discovery rate
HSCT: hematopoietic stem cell transplantation
HWE: Hardy-Weinberg equilibrium
KREC: kappa-deleting recombination excision circles
OLS: ordinary least squares
qPCR: quantitative polymerase chain reaction
ROC: receiver operating characteristic
RTE: recent thymic emigrant
SNP: single-nucleotide polymorphism
TREC: T-cell receptor excision circles

## References

1. Douek DC, McFarland RD, Keiser PH, Gage EA, Massey JM, Haynes BF, et al. Changes in thymic function with age and during the treatment of HIV infection. Nature (1998) 396:690–5. doi: 10.1038/25374

2. Poulin JF, Viswanathan MN, Harris JM, Komanduri KV, Wieder E, Ringuette N, et al. Direct evidence for thymic function in adult humans. J Exp Med (1999) 190:479–86. doi: 10.1084/jem.190.4.479

3. Hazenberg MD, Verschuren MC, Hamann D, Miedema F, van Dongen JJ. T cell receptor excision circles as markers for recent thymic emigrants: basic aspects, technical approach, and guidelines for interpretation. J Mol Med (Berl) (2001) 79:631–40. doi: 10.1007/s001090100271

4. Kimmig S, Przybylski GK, Schmidt CA, Laurisch K, Möwes B, Radbruch A, et al. Two subsets of naive T helper cells with distinct T cell receptor excision circle content in human adult peripheral blood. J Exp Med (2002) 195:789–94. doi: 10.1084/jem.20011756

5. Naylor K, Li G, Vallejo AN, Lee WW, Koetz K, Bryl E, et al. The influence of age on T cell generation and TCR diversity. J Immunol (2005) 174:7446–52. doi: 10.4049/jimmunol.174.11.7446

6. Mitchell WA, Meng I, Nicholson SA, Aspinall R. Thymic output, ageing and zinc. Biogerontology (2006) 7:461–70. doi: 10.1007/s10522-006-9061-7

7. Westermann J, Pabst R. Distribution of lymphocyte subsets and natural killer cells in the human body. Clin Investig (1992) 70:539–44. doi: 10.1007/BF00184787

8. Sottini A, Ghidini C, Zanotti C, Chiarini M, Caimi L, Lanfranchi A, et al. Simultaneous quantification of recent thymic T-cell and bone marrow B-cell emigrants in patients with primary immunodeficiency. Clin Immunol (2010) 136:217–27. doi: 10.1016/j.clim.2010.04.005

9. van Zelm MC, Szczepanski T, van der Burg M, van Dongen JJM. Replication history of B lymphocytes reveals homeostatic proliferation and extensive antigen-induced B cell expansion. J Exp Med (2007) 204:645–55. doi: 10.1084/jem.20060964

10. van Zelm MC, van der Burg M, Langerak AW, van Dongen JJM. PID comes full circle: applications of V(D)J recombination excision circles in research, diagnostics and newborn screening of primary immunodeficiency disorders. Front Immunol (2011) 2:12. doi: 10.3389/fimmu.2011.00012

11. Chan K, Puck JM. Development of population-based newborn screening for severe combined immunodeficiency. J Allergy Clin Immunol (2005) 115:391–8. doi: 10.1016/j.jaci.2004.10.012

12. Morinishi Y, Imai K, Nakagawa N, Sato H, Horiuchi K, Ohtsuka Y, et al. Identification of severe combined immunodeficiency by T-cell receptor excision circles quantification using neonatal Guthrie cards. J Pediatr (2009) 155:829–33. doi: 10.1016/j.jpeds.2009.05.026

13. Routes JM, Grossman WJ, Verbsky J, Laessig RH, Hoffman GL, Brokopp CD, et al. Statewide newborn screening for severe T-cell lymphopenia. JAMA (2009) 302:2465–70. doi: 10.1001/jama.2009.1806

14. Gerstel-Thompson JL, Wilkey JF, Baptiste JC, Navas JS, Pai SY, Pass KA, et al. High-throughput multiplexed T-cell-receptor excision circle quantitative PCR assay with internal controls for detection of severe combined immunodeficiency in population-based newborn screening. Clin Chem (2010) 56:1466–74. doi: 10.1373/clinchem.2010.144915

15. Nakagawa N, Imai K, Kanegane H, Sato H, Yamada M, Kondoh K, et al. Quantification of kappa-deleting recombination excision circles in Guthrie cards for the identification of early B-cell maturation defects. J Allergy Clin Immunol (2011) 128:223–5.e2. doi: 10.1016/j.jaci.2011.01.052

16. Borte S, von Döbeln U, Fasth A, Wang N, Janzi M, Winiarski J, et al. Neonatal screening for severe primary immunodeficiency diseases using high-throughput triplex real-time PCR. Blood (2012) 119:2552–5. doi: 10.1182/blood-2011-08-371021

17. Verbsky JW, Baker MW, Grossman WJ, Hintermeyer M, Dasu T, Bonacci B, et al. Newborn screening for severe combined immunodeficiency; the Wisconsin experience (2008-2011). J Clin Immunol (2012) 32:82–8. doi: 10.1007/s10875-011-9609-4

18. Kwan A, Abraham RS, Currier R, Brower A, Andruszewski K, Abbott JK, et al. Newborn screening for severe combined immunodeficiency in 11 screening programs in the United States. JAMA (2014) 312:729–38. doi: 10.1001/jama.2014.9132

19. Kwan A, Puck JM. History and current status of newborn screening for severe combined immunodeficiency. Semin Perinatol (2015) 39:194–205. doi: 10.1053/j.semperi.2015.03.004

20. Barbaro M, Ohlsson A, Borte S, Jonsson S, Zetterström RH, King J, et al. Newborn screening for severe primary immunodeficiency diseases in Sweden - a 2-year pilot TREC and KREC screening study. J Clin Immunol (2017) 37:51–60. doi: 10.1007/s10875-016-0347-5

21. Puck JM. Newborn screening for severe combined immunodeficiency and T-cell lymphopenia. Immunol Rev (2019) 287:241–52. doi: 10.1111/imr.12729

22. Gordukova MA, Oskorbin IP, Mishukova OV, Zimin SB, Zinovieva NV, Davydova NV, et al. Development of a reagent kit for quantitative determination of TREC and KREC DNA molecules in whole blood and dried blood spots by multiplex real-time PCR. Med Immunol (Russia) (2015) 17:467–78. doi: 10.15789/1563-0625-2015-5-467-478

23. Steinmann GG, Klaus B, Müller-Hermelink HK. The involution of the ageing human thymic epithelium is independent of puberty. A morphometric study. Scand J Immunol (1985) 22:563–75. doi: 10.1111/j.1365-3083.1985.tb01916.x

24. Franceschi C, Bonafè M, Valensin S, Olivieri F, De Luca M, Ottaviani E, et al. Inflamm-aging: an evolutionary perspective on immunosenescence. Ann N Y Acad Sci (2000) 908:244–54. doi: 10.1111/j.1749-6632.2000.tb06651.x

25. Aw D, Silva AB, Palmer DB. Immunosenescence: emerging challenges for an ageing population. Immunology (2007) 120:435–46. doi: 10.1111/j.1365-2567.2007.02555.x

26. Goronzy JJ, Weyand CM. Mechanisms underlying T cell ageing. Nat Rev Immunol (2019) 19:573–83. doi: 10.1038/s41577-019-0180-1

27. Nikolich-Zugich J. The twilight of immunity: emerging concepts in aging of the immune system. Nat Immunol (2018) 19:10–9. doi: 10.1038/s41590-017-0006-x

28. Lopez-Otin C, Blasco MA, Partridge L, Serrano M, Kroemer G. The hallmarks of aging. Cell (2013) 153:1194–217. doi: 10.1016/j.cell.2013.05.039

29. Lopez-Otin C, Blasco MA, Partridge L, Serrano M, Kroemer G. Hallmarks of aging: an expanding universe. Cell (2023) 186:243–78. doi: 10.1016/j.cell.2022.11.001

30. Montecino-Rodriguez E, Berent-Maoz B, Dorshkind K. Causes, consequences, and reversal of immune system aging. J Clin Invest (2013) 123:958–65. doi: 10.1172/JCI64096

31. Palmer DB. The effect of age on thymic function. Front Immunol (2013) 4:316. doi: 10.3389/fimmu.2013.00316

32. Lynch HE, Goldberg GL, Chidgey A, Van den Brink MRM, Boyd R, Sempowski GD. Thymic involution and immune reconstitution. Trends Immunol (2009) 30:366–73. doi: 10.1016/j.it.2009.04.003

33. Britanova OV, Putintseva EV, Shugay M, Merzlyak EM, Turchaninova MA, Staroverov DB, et al. Age-related decrease in TCR repertoire diversity measured with deep and normalized sequence profiling. J Immunol (2014) 192:2689–98. doi: 10.4049/jimmunol.1302064

34. Appay V, Sauce D. Naive T cells: the crux of cellular immune aging? Exp Gerontol (2014) 54:90–3. doi: 10.1016/j.exger.2014.01.003

35. Westera L, van Hoeven V, Drylewicz J, Spierenburg G, van Velzen JF, de Boer RJ, et al. Lymphocyte maintenance during healthy aging requires no substantial alterations in cellular turnover. Aging Cell (2015) 14:219–27. doi: 10.1111/acel.12311

36. Khan N, Shariff N, Cobbold M, Bruton R, Ainsworth JA, Sinclair AJ, et al. Cytomegalovirus seropositivity drives the CD8 T cell repertoire toward greater clonality in healthy elderly individuals. J Immunol (2002) 169:1984–92. doi: 10.4049/jimmunol.169.4.1984

37. Sylwester AW, Mitchell BL, Edgar JB, Taormina C, Pelte C, Ruchti F, et al. Broadly targeted human cytomegalovirus-specific CD4+ and CD8+ T cells dominate the memory compartments of exposed subjects. J Exp Med (2005) 202:673–85. doi: 10.1084/jem.20050882

38. Derhovanessian E, Maier AB, Hähnel K, Beck R, de Craen AJM, Slagboom EP, et al. Infection with cytomegalovirus but not herpes simplex virus induces the accumulation of late-differentiated CD4+ and CD8+ T-cells in humans. J Gen Virol (2011) 92:2746–56. doi: 10.1099/vir.0.036004-0

39. Wertheimer AM, Bennett MS, Park B, Uhrlaub JL, Martinez C, Pulko V, et al. Aging and cytomegalovirus infection differentially and jointly affect distinct circulating T cell subsets in humans. J Immunol (2014) 192:2143–55. doi: 10.4049/jimmunol.1301721

40. Allman D, Miller JP. B cell development and receptor diversity during aging. Curr Opin Immunol (2005) 17:463–7. doi: 10.1016/j.coi.2005.07.002

41. Cancro MP, Hao Y, Scholz JL, Riley RL, Frasca D, Dunn-Walters DK, et al. B cells and aging: molecules and mechanisms. Trends Immunol (2009) 30:313–8. doi: 10.1016/j.it.2009.04.005

42. Frasca D, Blomberg BB. Aging affects human B cell responses. J Clin Immunol (2011) 31:430–5. doi: 10.1007/s10875-010-9501-7

43. Olsen NJ, Kovacs WJ. Gonadal steroids and immunity. Endocr Rev (1996) 17:369–84. doi: 10.1210/edrv-17-4-369

44. Sutherland JS, Goldberg GL, Hammett MV, Uldrich AP, Berzins SP, Heng TSP, et al. Activation of thymic regeneration in mice and humans following androgen blockade. J Immunol (2005) 175:2741–53. doi: 10.4049/jimmunol.175.4.2741

45. Goldberg GL, Sutherland JS, Hammett MV, Milton MK, Heng TSP, Chidgey AP, et al. Sex steroid ablation enhances immune reconstitution following allogeneic hemopoietic stem cell transplantation. J Immunol (2007) 178:7473–84. doi: 10.4049/jimmunol.178.11.7473

46. Sakiani S, Olsen NJ, Kovacs WJ. Gonadal steroids and humoral immunity. Nat Rev Endocrinol (2013) 9:56–62. doi: 10.1038/nrendo.2012.206

47. Velardi E, Tsai JJ, Holland AM, Wertheimer T, Yu VWC, Zakrzewski JL, et al. Sex steroid blockade enhances thymopoiesis by modulating Notch signaling. J Exp Med (2014) 211:2341–9. doi: 10.1084/jem.20131289

48. Gellert M. V(D)J recombination: RAG proteins, repair factors, and regulation. Annu Rev Biochem (2002) 71:101–32. doi: 10.1146/annurev.biochem.71.090501.150203

49. Krangel MS. Mechanics of T cell receptor gene rearrangement. Curr Opin Immunol (2009) 21:133–9. doi: 10.1016/j.coi.2009.03.009

50. Shih HY, Krangel MS. Distinct contracted conformations of the Tcra/Tcrd locus during Tcra and Tcrd recombination. J Exp Med (2010) 207:1835–41. doi: 10.1084/jem.20100772

51. Schatz DG, Swanson PC. V(D)J recombination: mechanisms of initiation. Annu Rev Genet (2011) 45:167–202. doi: 10.1146/annurev-genet-110410-132552

52. Chen L, Carico Z, Shih HY, Krangel MS. A discrete chromatin loop in the mouse Tcra-Tcrd locus shapes the TCRdelta and TCRalpha repertoires. Nat Immunol (2015) 16:1085–93. doi: 10.1038/ni.3232

53. Hernandez-Munain C. Recent insights into the transcriptional control of the Tcra/Tcrd locus by distant enhancers during the development of T-lymphocytes. Transcription (2015) 6:65–73. doi: 10.1080/21541264.2015.1078429

54. Carico ZM, Roy Choudhury K, Zhang B, Zhuang Y, Krangel MS. Tcrd rearrangement redirects a processive Tcra recombination program to expand the Tcra repertoire. Cell Rep (2017) 19:2157–73. doi: 10.1016/j.celrep.2017.05.045

55. Orru V, Steri M, Sole G, Sidore C, Virdis F, Dei M, et al. Genetic variants regulating immune cell levels in health and disease. Cell (2013) 155:242–56. doi: 10.1016/j.cell.2013.08.041

56. Roederer M, Quaye L, Mangino M, Beddall MH, Mahnke Y, Chattopadhyay P, et al. The genetic architecture of the human immune system: a bioresource for autoimmunity and disease pathogenesis. Cell (2015) 161:387–403. doi: 10.1016/j.cell.2015.02.046

57. Brodin P, Jojic V, Gao T, Bhattacharya S, Angel CJL, Furman D, et al. Variation in the human immune system is largely driven by non-heritable influences. Cell (2015) 160:37–47. doi: 10.1016/j.cell.2014.12.020

58. Clave E, Araujo IL, Alanio C, Patin E, Bergstedt J, Urrutia A, et al. Human thymopoiesis is influenced by a common genetic variant within the TCRA-TCRD locus. Sci Transl Med (2018) 10:eaao2966. doi: 10.1126/scitranslmed.aao2966

59. Krenger W, Blazar BR, Holländer GA. Thymic T-cell development in allogeneic stem cell transplantation. Blood (2011) 117:6768–77. doi: 10.1182/blood-2011-02-334623

60. Gaballa A, Norberg A, Stikvoort A, Mattsson J, Sundberg B, Uzunel M, et al. Assessment of TREC, KREC and telomere length in long-term survivors after allogeneic HSCT: the role of GvHD and graft source and evidence for telomere homeostasis in young recipients. Bone Marrow Transplant (2018) 53:69–77. doi: 10.1038/bmt.2017.216

61. Tsamadou C, Gowdavally S, Platzbecker U, Sala E, Valerius T, Wagner-Drouet E, et al. Donor genetic determinant of thymopoiesis rs2204985 impacts clinical outcome after single HLA mismatched hematopoietic stem cell transplantation. Bone Marrow Transplant (2022) 57:1539–47. doi: 10.1038/s41409-022-01751-1

62. Tobin J. Estimation of relationships for limited dependent variables. Econometrica (1958) 26:24–36. doi: 10.2307/1907382

63. Cragg JG. Some statistical models for limited dependent variables with application to the demand for durable goods. Econometrica (1971) 39:829–44. doi: 10.2307/1909582

64. Mullahy J. Specification and testing of some modified count data models. J Econom (1986) 33:341–65. doi: 10.1016/0304-4076(86)90002-3

65. Koenker R, Bassett G. Regression quantiles. Econometrica (1978) 46:33–50. doi: 10.2307/1913643

66. Efron B, Tibshirani RJ. An Introduction to the Bootstrap. New York: Chapman & Hall (1993). 436 p.

67. DeLong ER, DeLong DM, Clarke-Pearson DL. Comparing the areas under two or more correlated receiver operating characteristic curves: a nonparametric approach. Biometrics (1988) 44:837–45. doi: 10.2307/2531595

68. Benjamini Y, Hochberg Y. Controlling the false discovery rate: a practical and powerful approach to multiple testing. J R Stat Soc Series B Stat Methodol (1995) 57:289–300. doi: 10.1111/j.2517-6161.1995.tb02031.x

69. Duan N. Smearing estimate: a nonparametric retransformation method. J Am Stat Assoc (1983) 78:605–10. doi: 10.1080/01621459.1983.10478017

70. Wilson EB. Probable inference, the law of succession, and statistical inference. J Am Stat Assoc (1927) 22:209–12. doi: 10.1080/01621459.1927.10502953

